# Development of the foregut in *Katharina tunicata* (Mollusca; Polyplacophora)

**DOI:** 10.1101/2021.02.28.433208

**Authors:** Brandy S. Biggar

## Abstract

As a highly diverse phyla, Mollusca is both intriguing to study and difficult to define. No single common feature distinguishes the phyla, but the most common and recognizable is the radula. Throughout Molluscan history, there have been many developments to the radular structure. Evolutionary development of unique radular structures may have been made possible by developmental modularity. This histological study examines the development of polyplacophoran internal gut development. Here, I describe the developmental sequence of chiton feeding structures for the first time. Feeding structures are present by 10 days post hatch. Future research should examine larval stages prior to this in order to determine whether developmental modularity exists in polyplacophorans.

## Introduction

With over 100,000 described species, Mollusca is one of the largest and most successful metazoan phyla (Seed, 1983). Although no single character defines the phylum, the radula is one of the most distinctive and easily identifiable features, and is present in members of all molluscan classes except Bivalvia (Runnegar and Pojeta, 1985; Salvini-Plawen, 1988; Trueman and Clarke, 1988; Purchon, 2013). Fossil records date Mollusca back ~500 My to the Cambrian, but the abrupt presence of 15 genera in the Phanerozoic records suggest a Precambrian presence that may yet await discovery (Runnegar and Pojeta, 1985; Trueman and Clarke, 1988; Todt et al., 2008). The fossil record includes trace fossils from Ediacaran strata, radula-like fossils from the mid-Cambrian, and *definitive* radular evidence from the Ordovician (Scheltema et al., 2003; Todt et al., 2008). The fossil record suggests that radular rasping may be the primitive molluscan feeding mode (Seed, 1983). The presence of this unique feeding structure throughout molluscan history is suggestive of its importance within the clade, and possible presence in the molluscan common ancestor.

The polyplacophoran radula is unique among molluscs encompassing key features from each of the other major types; these features allow them to simultaneously excavate and sweep food from the substratum (Steneck and Watling, 1982). With robust buccal muscles and biominerally hardened denticles, the chiton radula is capable of generating immense force (Steneck and Watling, 1982). Bilaterally symmetric ribbons of transparent proteinaceous material hold 25 to 150 rows of teeth in a lateral arrangement of 17 teeth per row, occupying up to 1/3 of the body length (Schwabe and Wehrtmann, 2009). Four teeth per row are used in grazing, two dominant and two marginal (Steneck and Watling, 1982). Dominant teeth are often tricuspid and heavily mineralized for sediment excavation; x-ray diffraction shows the presence of magnetite (an iron-containing biomineral), goethite, lepidocrocite, carbonate apatite, and magnetite occurs in varying degrees per species (Lowenstam, 1962; Steneck and Watling, 1982).

The ancestral life-history and evolution of the molluscan life cycle are still extensively debated. Extant molluscs’ broadly fall under two of the three main types of life-history patterns: planktonic feeding larvae (planktotrophic), and planktonic non-feeding larvae (lecithotrophic), with isolated instances of the third type (direct development). The two major hypotheses regarding life history evolution of the molluscan ancestor are the ‘larval-first’ hypothesis and the ‘intercalation’ hypothesis (Page, 2009). According to the larval-first hypothesis, or “Trochaea theory”, the ancestral molluscan condition was holoplanktonic. Later, these organisms adopted a benthic lifestyle later in development and postponed sexual maturation to the benthic stage. Eventually, the initial planktonic stage of the life history, which must have been capable of feeding, became the planktotrophic larvae of a life history that was otherwise mostly benthic. The intercalation theory starts with a holobenthic organism, which later acquiring a planktonic juvenile stage. Because the planktonic stage is added secondarily, structures for planktonic feeding did not exist and the larva was therefore lecithotrophic. Subsequent development of larval feeding is then a derived state under the intercalation hypothesis (Haszprunar et al., 1995).

Basal clades may preserve the ancestral state, so mapping life history on to phylogeny can help resolve the ancestral state and polarity of change for molluscan life history evolution. Within Gastropoda, Patellogastropoda and Vetigastropoda are generally considered the basal clades; within Mollusca, Polyplacophora is considered basal. All three of these clades produce lecithotrophic larvae. Furthermore, among extant gastropods, the smallest species almost always exhibit direct development or lecithotrophic larvae and the fossil record suggests that the first molluscs were very small (Chaffee and Lindberg, 1986). The notion that the first molluscs did not produce planktotrophic lavae is also corroborated by the fossil record (Nützel et al., 2006). Finally, comparative studies of cleavage patterns during early development of spiralians have also been interpreted as supporting the notion of ancestral lecithotrophy rather than planktotrophy (Guralnick and Lindberg, 2001). Clades of gastropods with planktotrophic larvae show the largest diversity and complexity in feeding systems (Page and Hookham, 2017). If complex feeding systems are the derived state, then planktotrophy may enhance evolvability of foregut development.

Resolving the molluscan phylogeny has been regarded as “one of the greatest challenges in invertebrate evolution” (Todt et al., 2008; Sigwart et al., 2013; Schrödl and Stöger, 2014). The two major competing hypotheses are Aculifera/ Conchifera and Testaria/ Conchifera; both with Polyplacophora as a basal clade (Schrödl and Stöger, 2014). Many authors consider chitons to possess ancestral traits, making them an attractive study subject; yet little work has been done on chiton development. The majority of chiton research has focused on muscle, neuron, and eye development, while few have examined foregut development, and none have followed foregut development through metamorphosis. Here, I examine and describe the relative rate of foregut development in *Katharina tunicata*, a polyplacophoran mollusc, to provide insight into the basal mollusc as the possible ancestral condition.

## Methods

Larvae were reared and fixed prior to this study.

### Sectioning

Serial 1 μm thick sections were obtained using a manual rotary MT5000 Sorvall Ultra microtome. One cross-sectional series was prepared of a ten days-post-hatch (dph) specimen; all other sections were longitudinal. Four longitudinal section series were completed on larvae at the following stages: 10 dph, 13 dph, 17 dph (X2); another four were juvenile specimens: 1 day post-metamorphosis (dpm), 14 dpm, 35 dpm (X2). Sections were stained using Richardson’s stain and mounted on glass slides (Richardson et al., 1960).

### Describing development

Photomicrographs were obtained using a QImaging Retiga 2000R digital camera mounted on a Zeiss Axioskop microscope, operated by QCapture Pro 6.0 software. Contrast, brightness, and sharpness of images were adjusted using Adobe PhotoShop CS Version 19.1.3. Developmental events were reconstructed by examining photographs of sections through successive stages of development.

## Results

### Larval development

Hatched larvae were egg-shaped, measuring 290 μm long by 230 μm wide (Fig. 2). Larvae hatched with 125 μm cilia extending anteriorly (the apical tuft), surrounded by a collar of short 15 μm long microvilli, and a locomotory band of 70 μm long cilia encircling the body at its greatest girth (prototroch). As observed by Watanabe and Cox (1974), the larvae also had a short tuft of cilia (30 μm) protruding from the teleotrochal field at the posterior end of the body. At 2 dph, larvae had grown more linearly but were still egg-shaped with a wider pretrochal region and a tapered postrochal region (Fig. 3). Length was 340 μm and max width was 250 μm. The entire larval body was highly opaque.

At 4 dph, larval eyespots (diameter = 15 μm) and mouth were visible postrochally (Fig. 4). The larvae did not exhibit much growth in body size at this stage (360 μm × 250 μm), but the body appeared more barrel-shaped with the pre- and postrochal regions approaching similar width. The larva also appeared to have a small prototrochal bulge (~15 μm). Although the ventral side was still opaque and dull, the dorsal surface was beginning to reflect light, and the periphery was semi-transparent. A fine covering of microvilli extended from the rostral tip, across the ventral surface, and around the caudal tip; microvilli did not extend across the mantle field (dorsal side from caudal tip to just above prototroch; Fig. 5; Leise, 1984).

By 6 dph, the larvae had elongated, giving it a narrower profile with a near pill-shaped form. A pit was observed on the anterior tip where the apical tuft extended from the surface. The apical tuft and prototroch both remained near hatching length (140 μm, 75 μm respectively). The shell gland anlagen were beginning to take form on the dorsal surface as narrow ridges and grooves. Larval eyespots were observed, and the girdle surrounding the periphery was becoming nearly transparent.

Spicules and six shell valves were visible by 8 dph as thin mineral deposits. Five to six rows of spiniferous cells and their associated intracellular spicules could be seen lining the mantle field. The larval eyes were situated directly below (ventrally) the spicule band and behind (caudally) the prototroch (Fig. 6; Henry et al., 2004; Leise, 1984). At this point, large cells appear on the dorsal surface, particularly in the shell gland anlagen and the foot begins delineation (Fig. 7, 8).

The first larval stage that was sectioned, 10 dph, was observed to have eyespots (pigment only), buccal mass, radular teeth, radular cartilages, esophagus, neuropil, and spicules (Fig. 8). The mantle field spicules could be seen in a linear arrangement running dorsocaudally in the postrochal region, and a few pretrochally on the dorsal edge (Fig. 9; (Henry et al., 2004; Leise, 1984). In whole mounts, the foot could be seen along the ventral side as a large v-shaped shadowy structure (Fig. 10). By this stage, the prototroch was visible in greater detail and I could view the double layer of circumferential trochoblasts (Fig. 11).

Longitudinal sections through the flanks of 13 dph specimens showed large, clearly visible eyespots with pigment surrounding a colourless center (Fig. 12). By 13 dph the stomodeum and radular cartilages were much more apparent and differentiated (Fig. 13).

By 17 dph the larvae were elongated and dorsoventrally flattened with translucent periphery and highly delineated foot. The radular teeth could be seen extruding from the buccal mass as they were added to the radular ribbon (Fig. 14). At this stage the larvae still had apical tuft and prototroch.

### Metamorphosis

Larvae of *K. tunicata* metamorphosis is indicated by loss of the prototroch and apical ciliary tuft. At 1 dpm the foot was clearly delineated by the pallial groove and seven shell valves were clearly evident and beginning to extend towards each other longitudinally (Fig. 15, 16A, B). The juveniles were near-round and pale with white shell valves and hairs surrounding the margin of the girdle (Leise, 1984). By 3 dpm, valves were 50 μm wide and juveniles were 700 μm long by 450 μm wide (Fig. 16C).

At 14 dpm, the dorsal side of the juveniles was completely covered by white shell valves (Fig. 16D). The radula was well developed with magnetite-capped denticles and the esophagus could be traced dorsally to the stomach and intestine (Fig. 17). Cuticle was also clearly visible proximal to shell valves and the foot had an abundance of pedal glands (Fig. 15B).

By 36 dpm, there were still only 7 valves present, but they were each significantly wider (75 – 100 μm), extending towards each other and beginning to articulate (Fig. 16F). The valves also appeared to be differentiating into their separate regions (tegmentum, articulamentum). By this stage the radular denticles and their magnetite caps were highly differentiated and the radular cartilages were arranged in a linear order as opposed to the spherical structure observed at earlier stages. The esophagus could easily be seen lined with cilia and extending dorsocaudally. The observed developmental sequence is summarized in Table 1 (Appendix II).

## Discussion

Major developmental stages were observed to occur in the same order as previous research (± 10 days; Table 2). The overall developmental sequence observed in *K. tunicata* was hatching, larval eyes, spicules, shell plates, settlement, and metamorphosis. This developmental list focuses mainly on external morphology and behaviour because most previous developmental studies on polyplacophorans have not provided information on internal events of morphogenesis. Although Leise (1984) examined spicule formation via histological sections, the same was accomplished here without sectioning (Fig. 6). I was, however, unable to pinpoint the exact timing and location of spicule formation, unlike Watanabe and Cox (1974) who note that spicule formation began first in the pretrochal region.

This study went beyond the external morphological development to examine the timing of foregut ontogenesis. By examining histological sections of *K. tunicata* at 10, 13, and 17 dph, I was able to determine that the buccal mass, radular teeth, radular cartilages, and esophagus had all begun formation by 10 dph (Fig. 10). By 13 dph the cartilages and buccal mass had a more distinct appearance, and by 17 dph the radula could be seen beginning to extrude from the buccal mass. Magnetite caps on the radular teeth, a unique chiton feature, was evident in juveniles at 36 days after metamorphosis. The development of these structures has not been previously described in Polyplacophora.

This study attempted to narrow the timing of foregut development but was unable to determine the start of formation. Because there was already evidence of foregut structures in 10 dph larvae, examination of earlier larval stages is imperative. Also, because eyespots were evident by 4 dph, histological sections as early, or earlier, than 4 dph would have been beneficial to examine. Future work should re-examine this problem at earlier stages of chiton development. Further support for the intercalation theory could be achieved by examining foregut development in other potentially basal clades such as the aplacophorans, Caudofoveata and Solenogastres (Todt et al., 2008).

## Conclusions

This study is the first to describe internal polyplacophoran larval development. I show that the buccal mass, radular teeth, radular cartilages, and esophagus all begin formation by 10 days post hatch. Furthermore, I show that the radula extrudes from the buccal mass by 17 days post hatch, and magnetite caps are formed by 36 days post metamorphosis.

## Appendix I Figures

**Figure 1.**
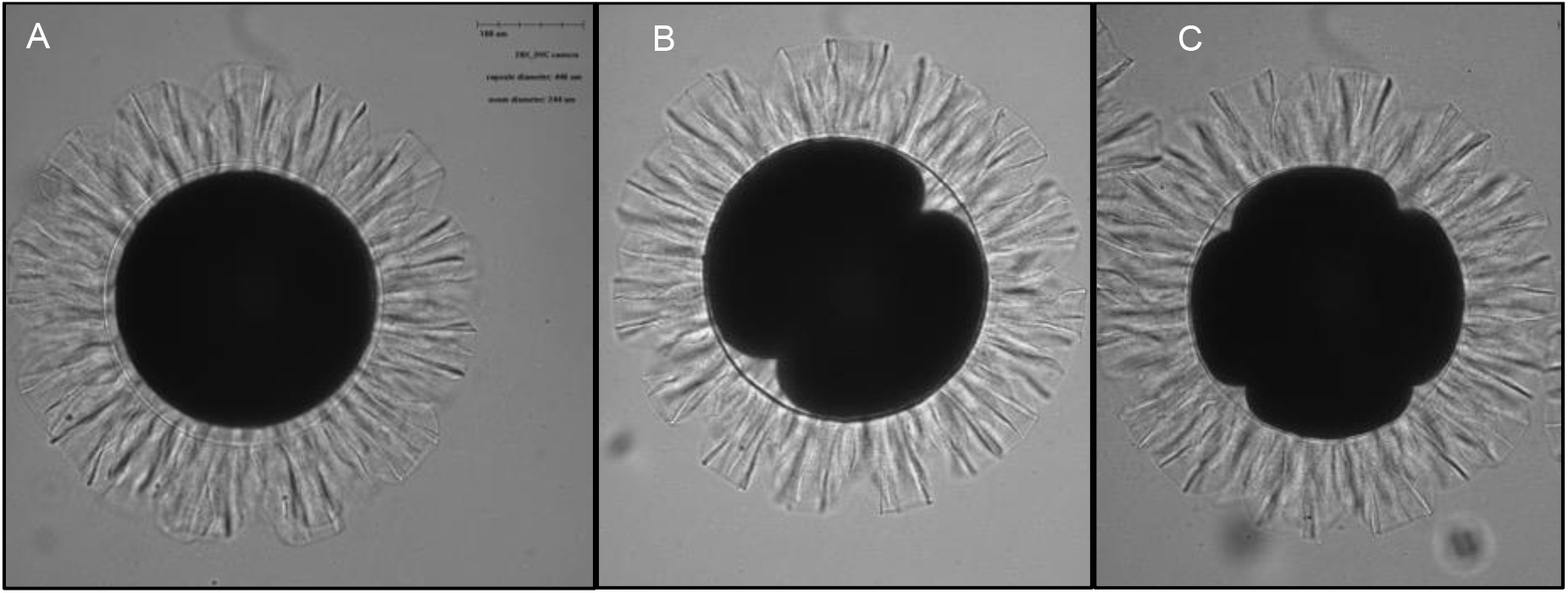
Early development of *Katharina tunicata* embryos with opaque yolky cytoplasm and transparent hull in A) zygote, B) 2-cell stage, and C) 4-cell stage.

**Figure 2.**
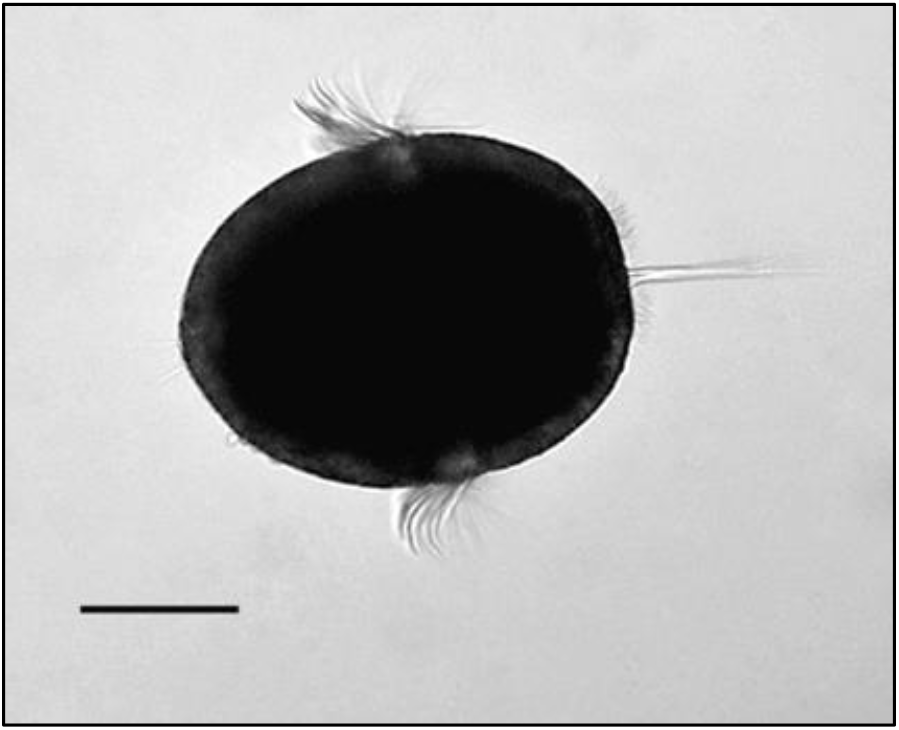
1 day post hatching larval *Katharina tunicata*, depicting clear apical tuft and prototroch. Scale bar represents 100μm.

**Figure 3.**
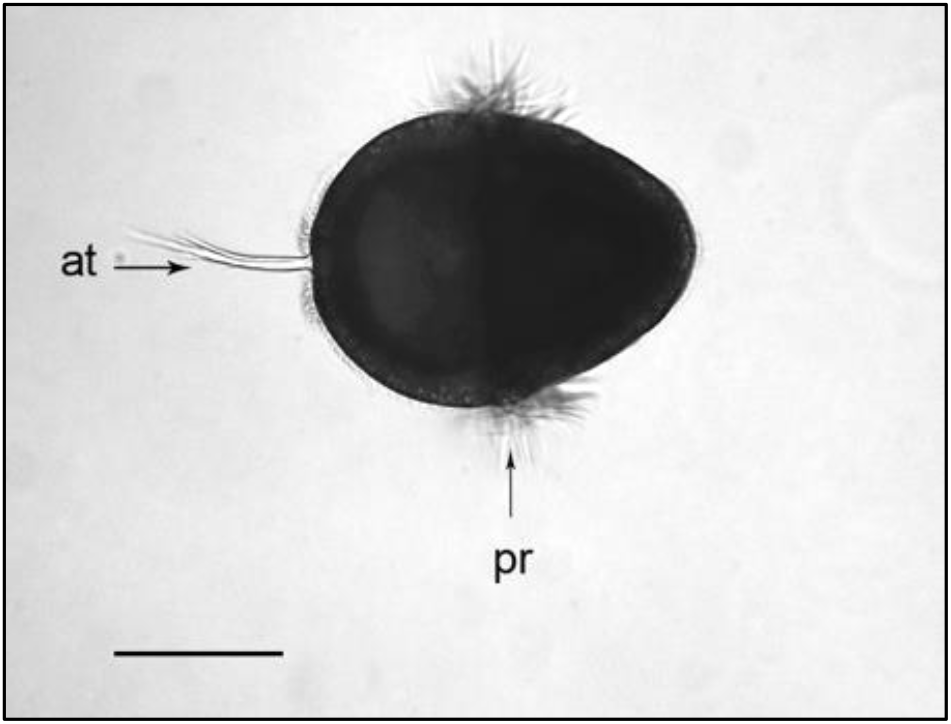
Egg-shaped 2 dph *K. tunicata* larvae with apical tuft surrounded by microvilli collar. Scale bar represents 150 μm; at, apical tuft; pr, prototroch.

**Figure 4.**
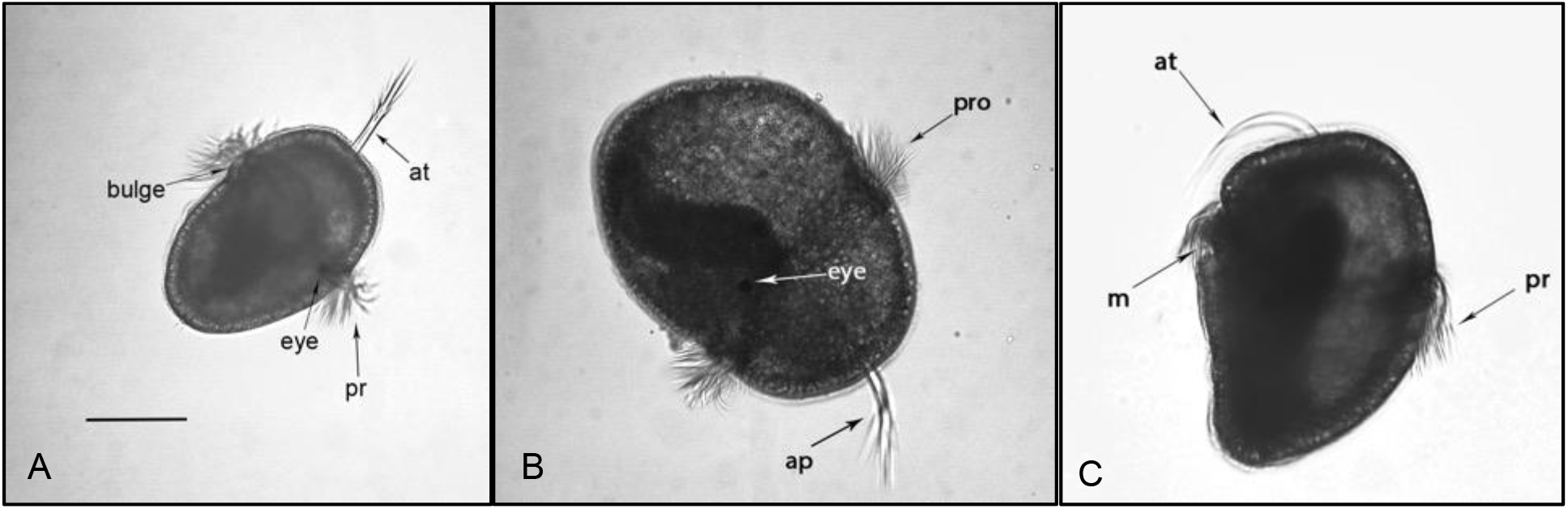
*Katharina tunicata* 4 dph larvae with eyespots, prototrochal bulge, and mouth. Scale bar represents 150 μm; at, apical tuft; m, mouth; pr, prototroch.

**Figure 5.**
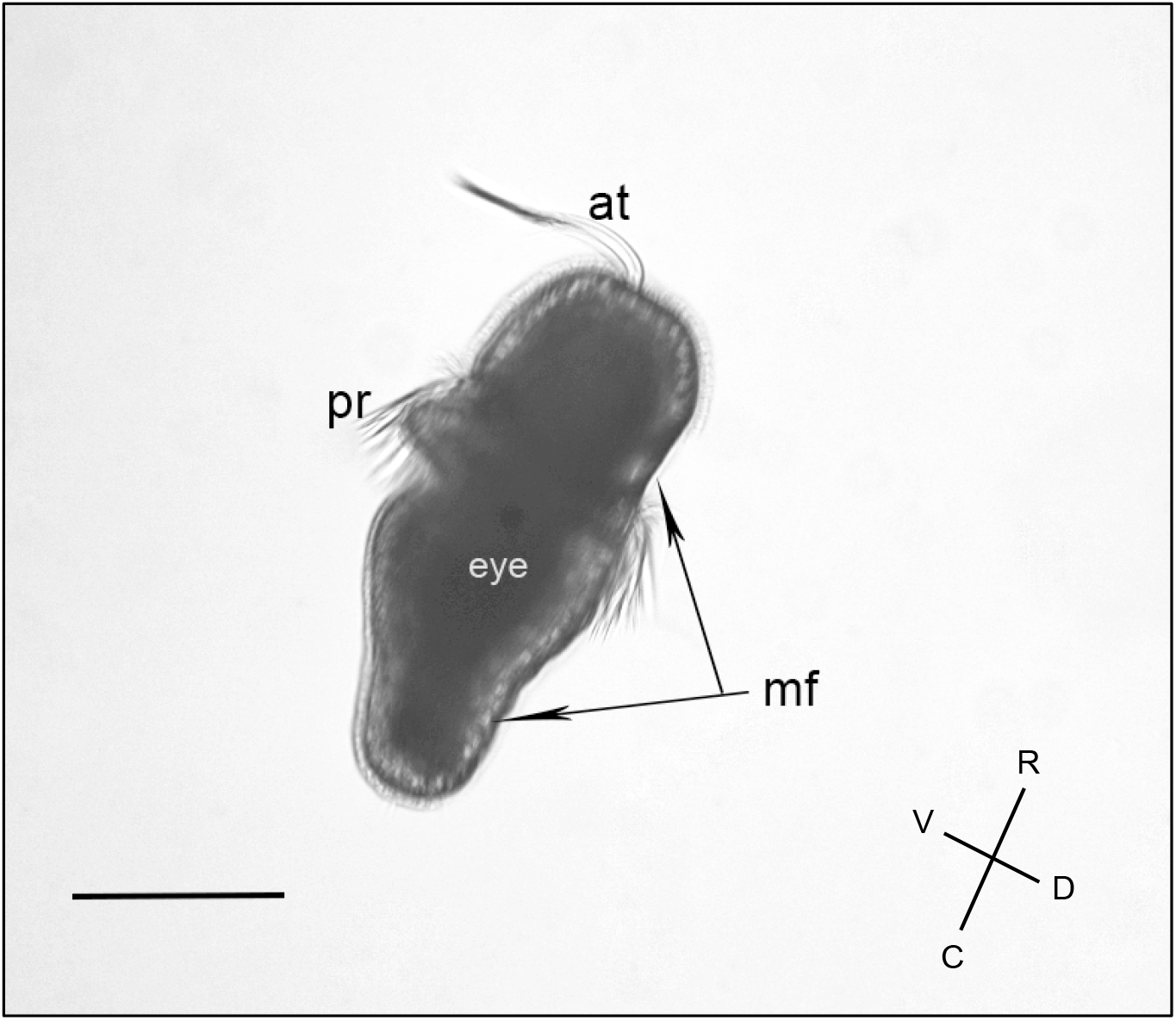
*Katharina tunicata* 6 dph larva displaying an already elongated body form. Mantle field is visible as a region devoid of microvilli on the dorsal side from the caudal end to mid-way up the pre-trochal region. Scale bar represents 150 μm and direction vector show the rostral, dorsal, ventral, and caudal axes of the larva. at, apical tuft; pr, prototroch; mf, mantle field.

**Figure 6.**
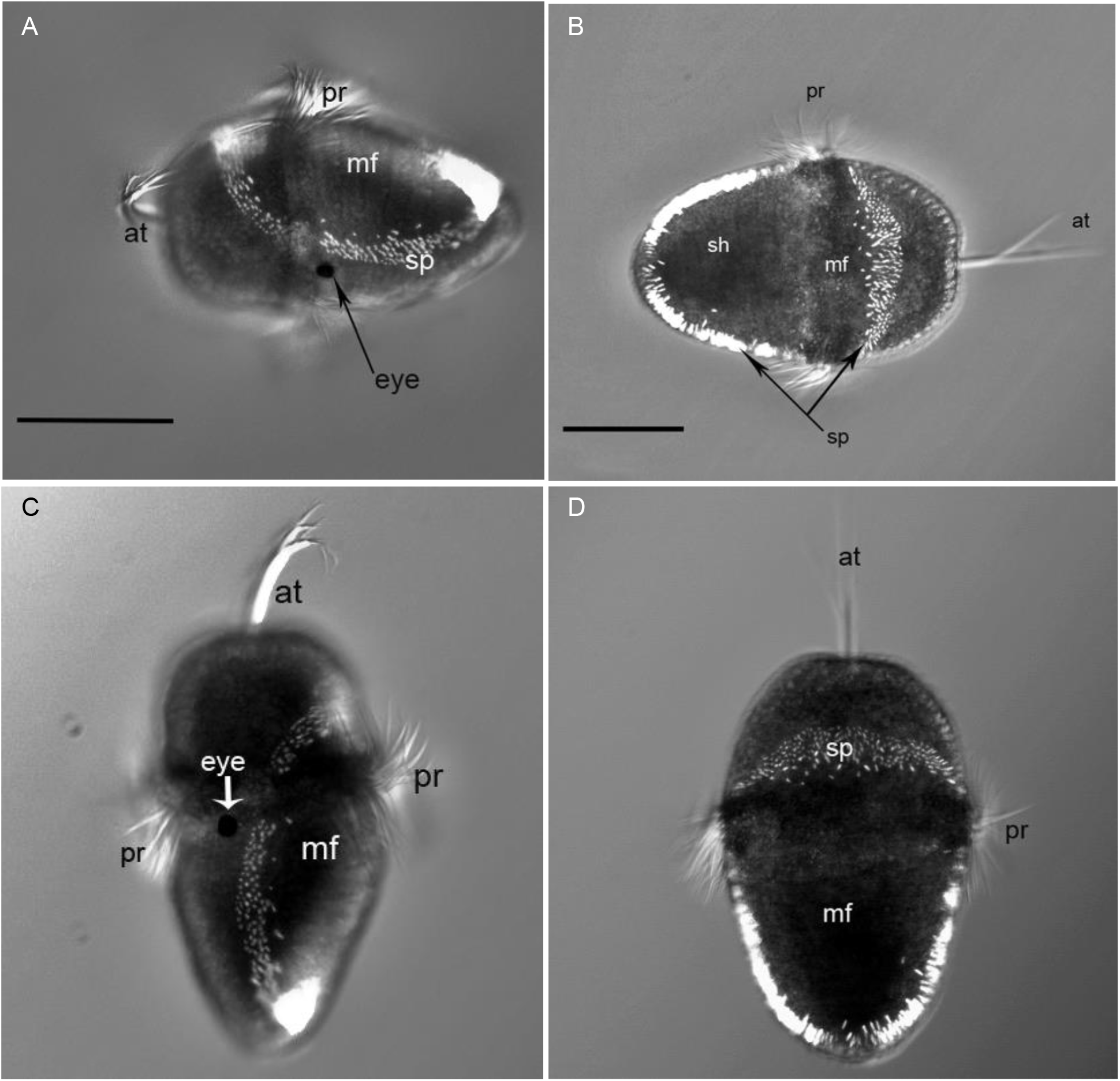
*Katharina tunicata* 8 dph larvae depicting the mantle field which is delineated by and encircling band of 5-6 rows of spicule-forming spiniferous cells. Scale bar represents 150 μm. at, apical tuft; pr, prototroch; mf, mantle field; sp, spicules; sh, shell anlagen.

**Figure 7.**
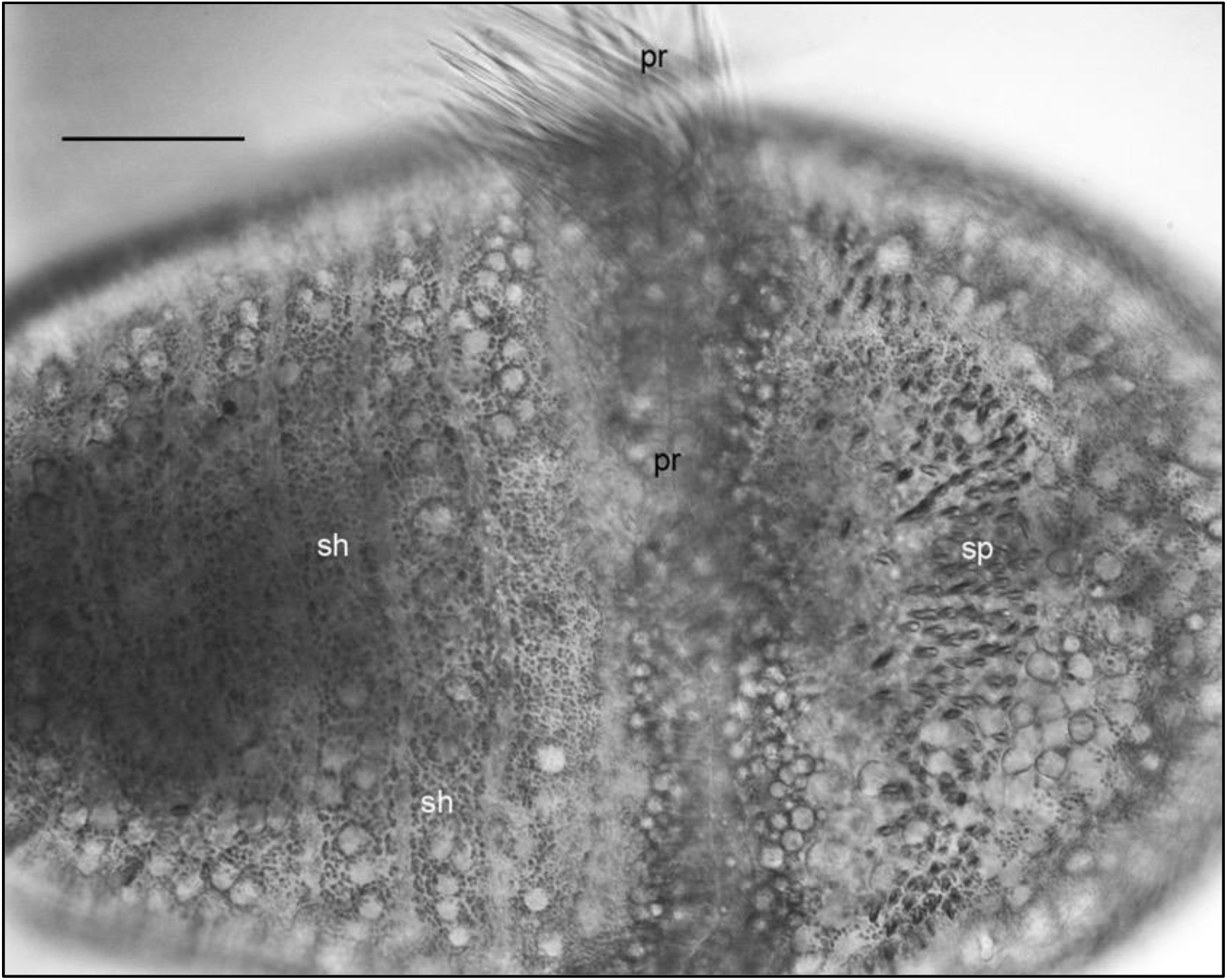
*Katharina tunicata* at 8 dph showing the delineated regions of shell anlagen postrochally along the mantle field, as well as pretrochal spicules, and two distinct rows of prototrochal cells. Scale bar is 50 μm; pr, prototroch; sh, shell anlage; sp, spicules.

**Figure 8.**
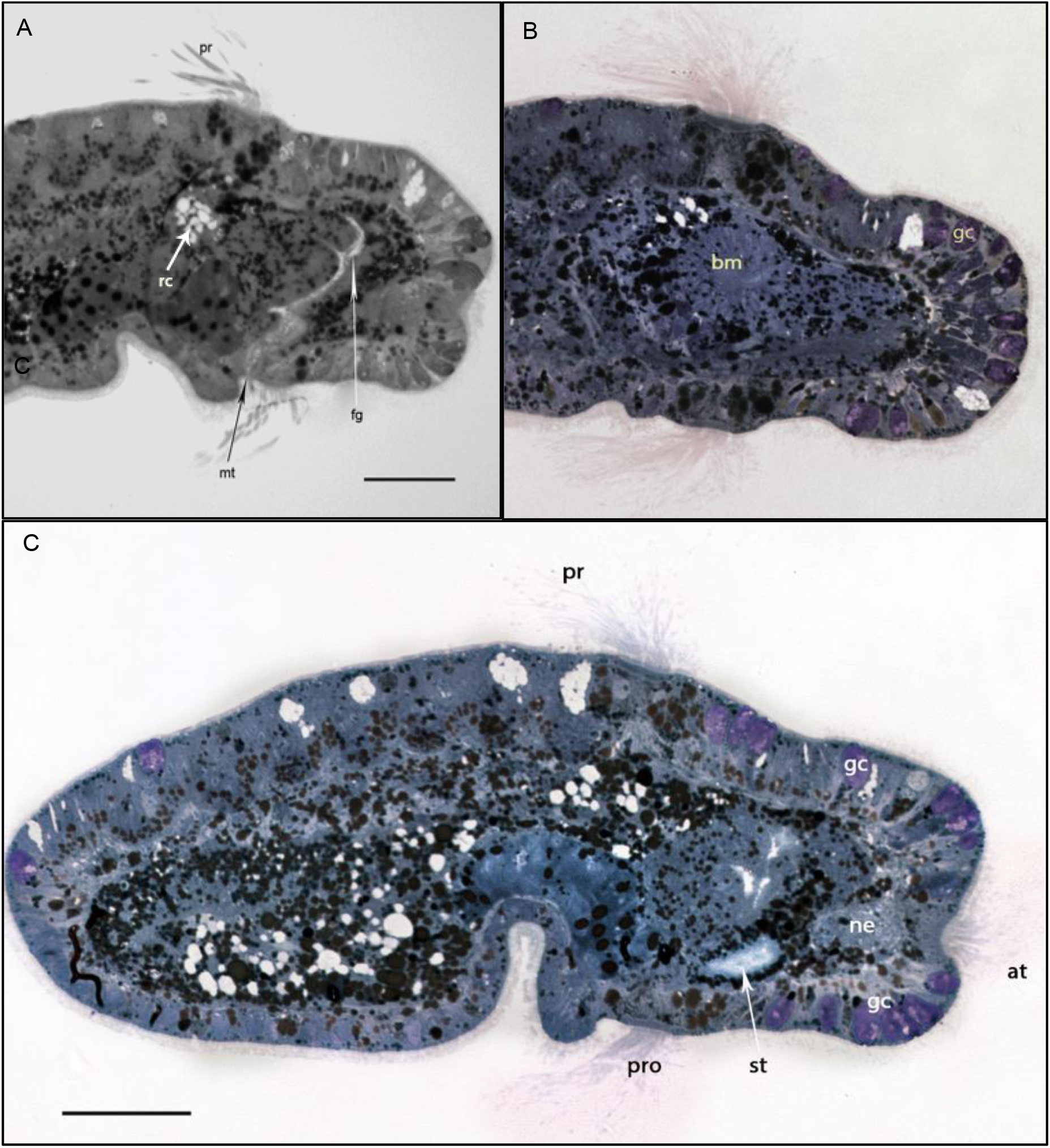
*Katharina tunicata* 10 days post hatch larvae 1 μm thickness histological sections displaying early developmental stages of foregut components. Scale bars are representative of 50 μm; pro, prototroch; gc, glandular cell; ne, neuropil; st, stomodeum; fg, foregut; mt, mouth.

**Figure 9.**
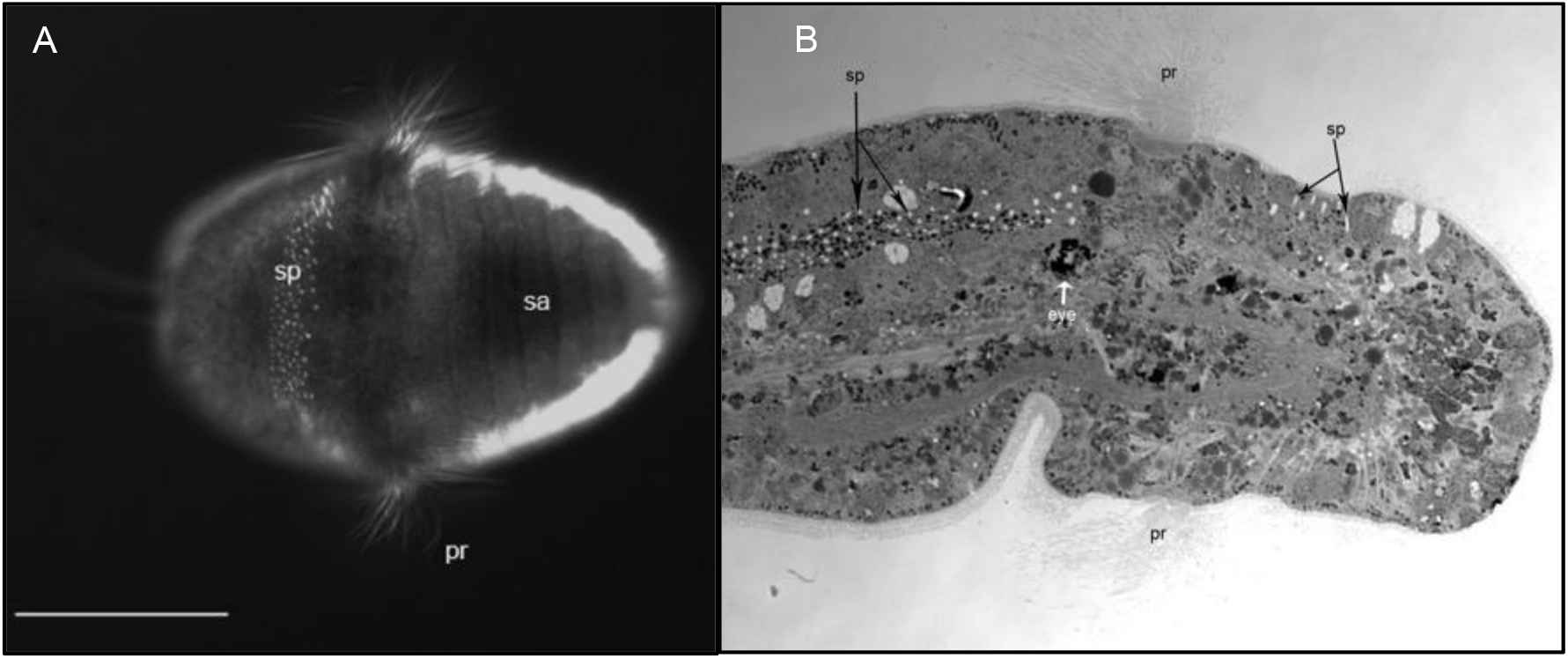
A) Dorsal view of *K. tunicata* 10 dph specimen displaying linear shell anlagen and spicules of the mantle field. B) Longitudinal histological section (1 μm thick) of *K. tunicata* (10 dph) larva. Scale bar represents 150 μm; pr, prototroch; sp, spicules; sa, shell anlage.

**Figure 10.**
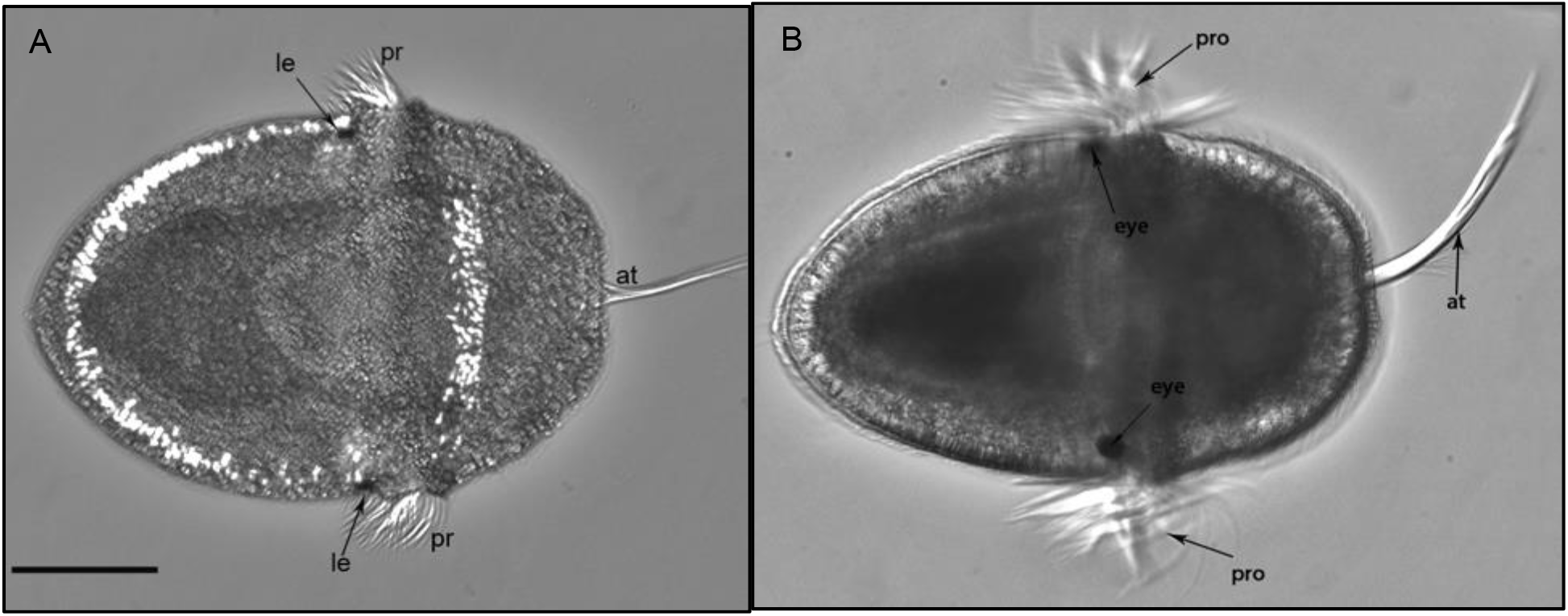
Whole specimen 10 dph *Katharina tunicata* larvae ventral view with postrochal larval eyes and foot visible. Scale bar represents 150 μm; at, apical tuft; le/eye, larval eye; pr/pro, prototroch.

**Figure 11.**
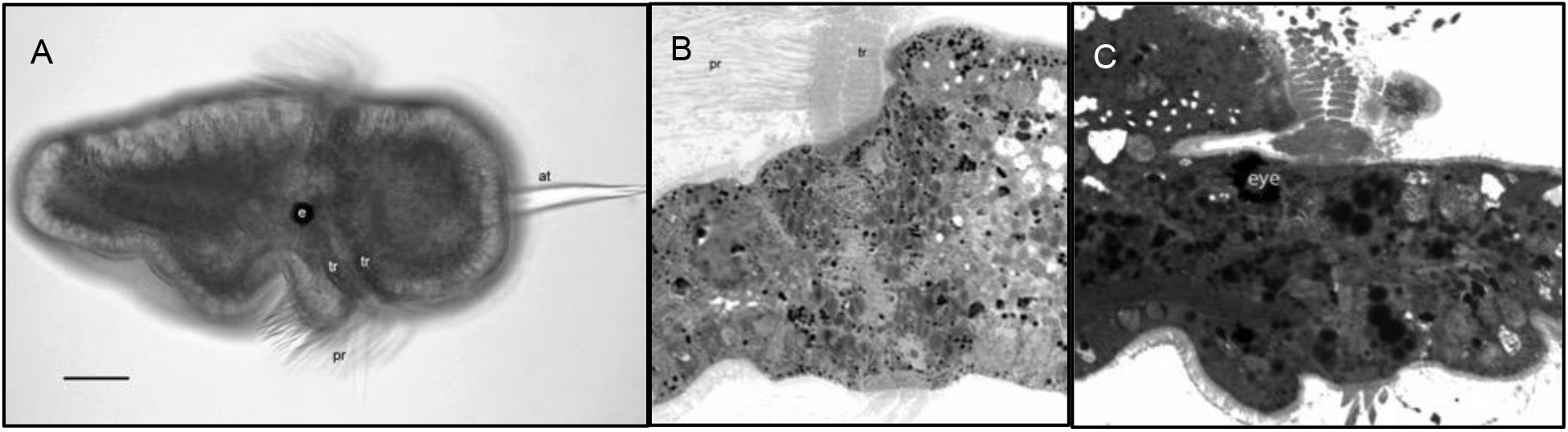
*Katharina tunicata* A) 10 dph whole mount and B, C) 17 dph 1 μm histological sections, showing a prototroch made up of two circumferential stacked rows of trochoblast cells. Scale bar represents 50 μm; pr, prototroch; tr, trochoblast; e/eye, larval eye; at, apical tuft.

**Figure 12.**
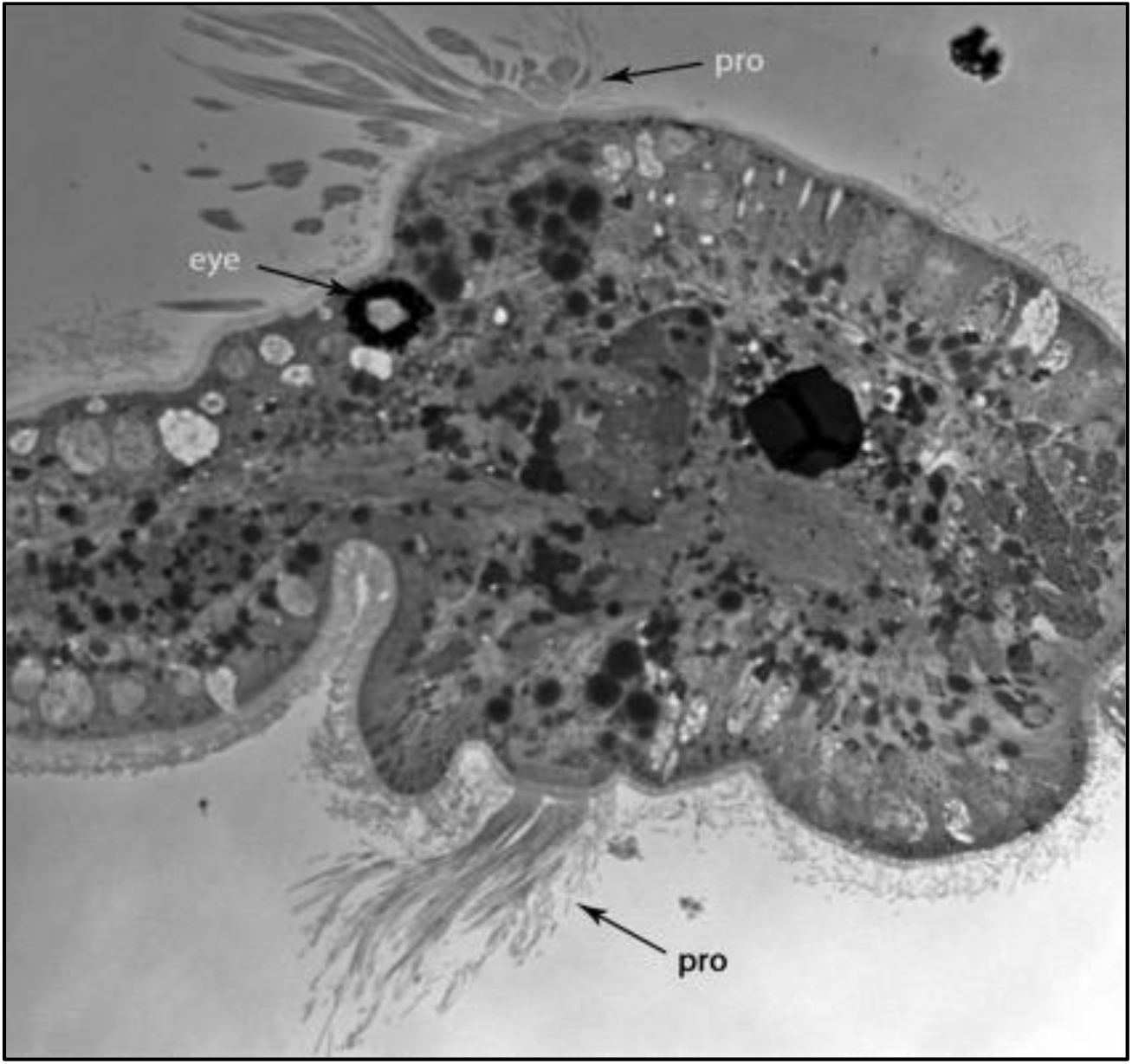
Transverse histological sections, 1 μm thick, of 13 dph *K. tunicata* larvae. Scale bar represents 50 μm; pro/pr, prototroch; ne, neuropil, at, apical tuft; mt, mouth; pg, pedal gland.

**Figure 13.**
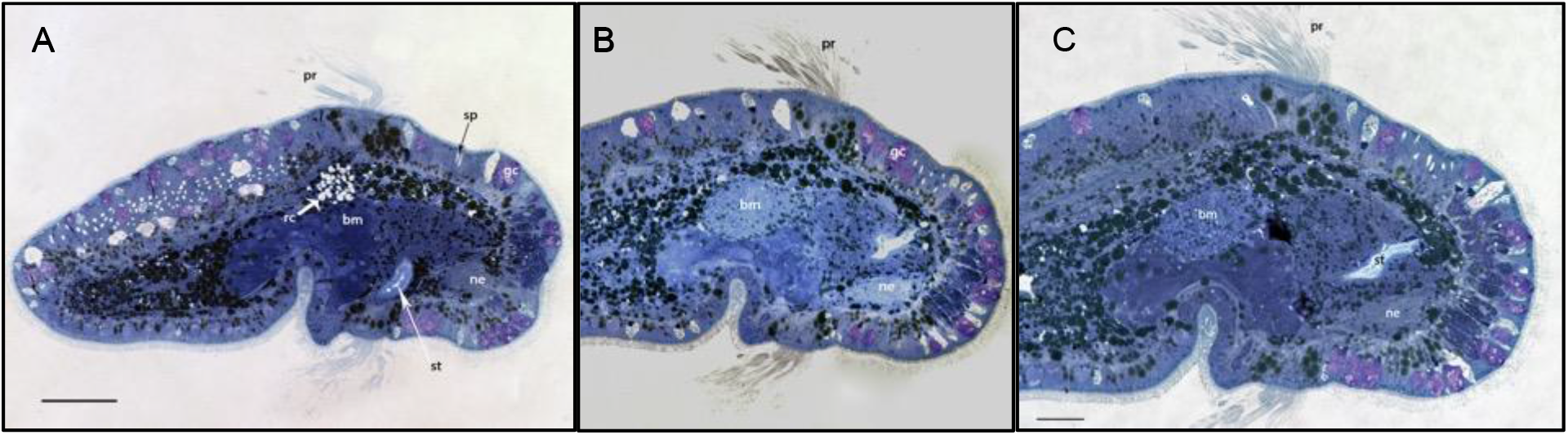
*K. tunicata* larvae, 1 μm, 13 dph midsagittal histological sections showing extent of foregut development. Scale bars represent 50 μm; bm, buccal mass; rc, radular cartilages; st, stomodeum; gc, glandular cell; pr, prototroch; sp, spicule; ne, neuropil.

**Figure 14.**
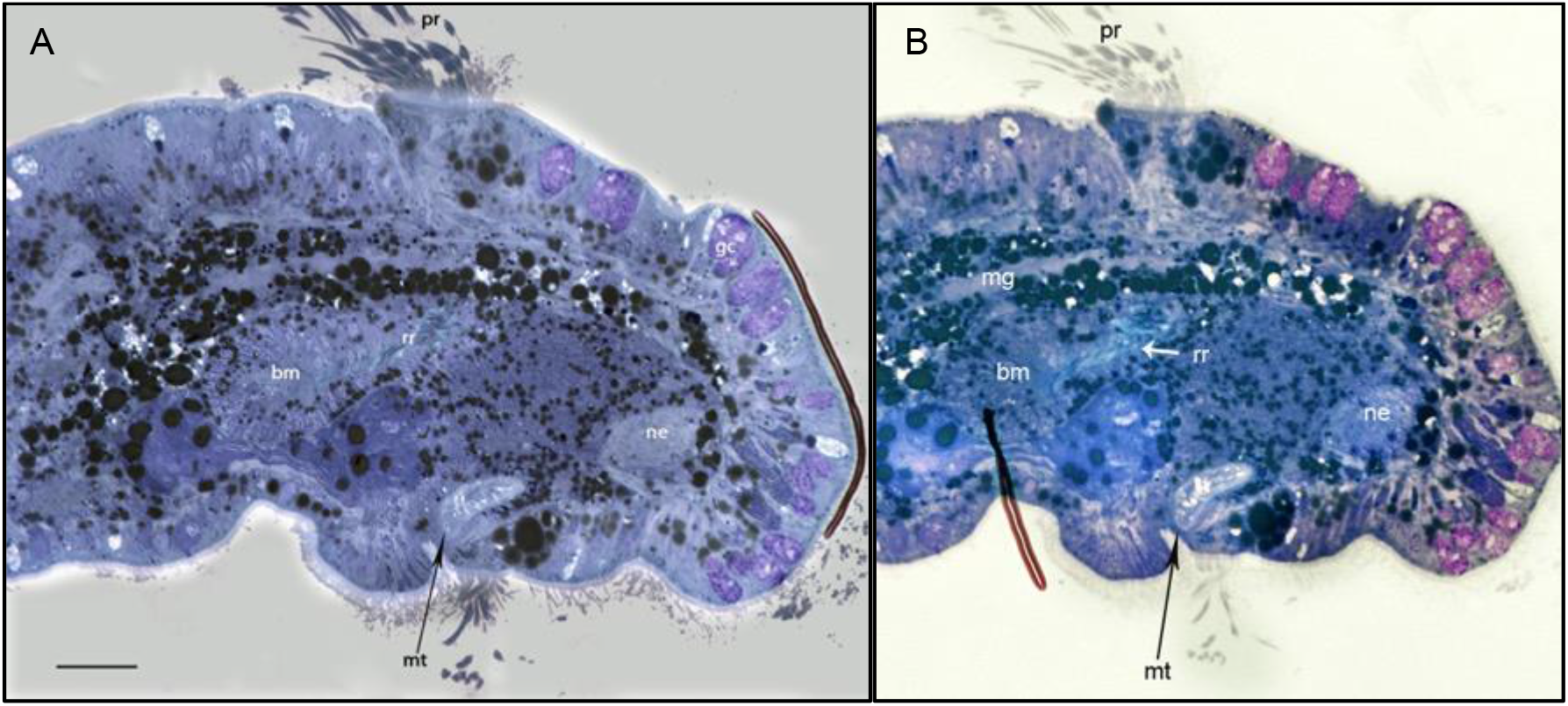
*Katharina tunicata* 17 dph larvae, 1 μm histological sections through mouth opening, with evidence of radular rudiment. Scale bar represents 50 μm; bm, buccal mass; pr, prototroch; mg, midgut; mt, mouth; ne, neuropil; rr, radular rudiment.

**Figure 15.**
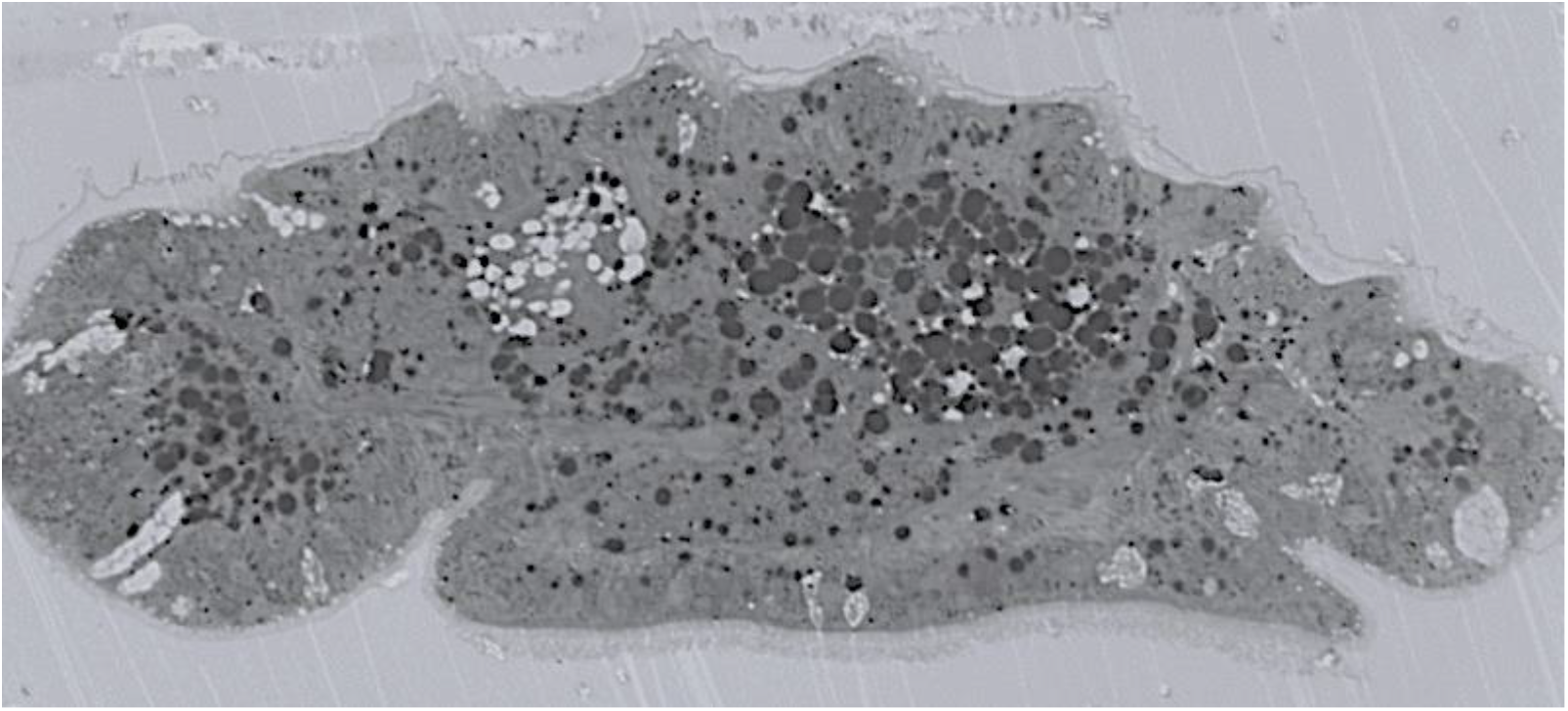
*K. tunicata* larva directly following metamorphosis (1 dpm) with a clear view of the delineated foot and pallial groove.

**Figure 16.**
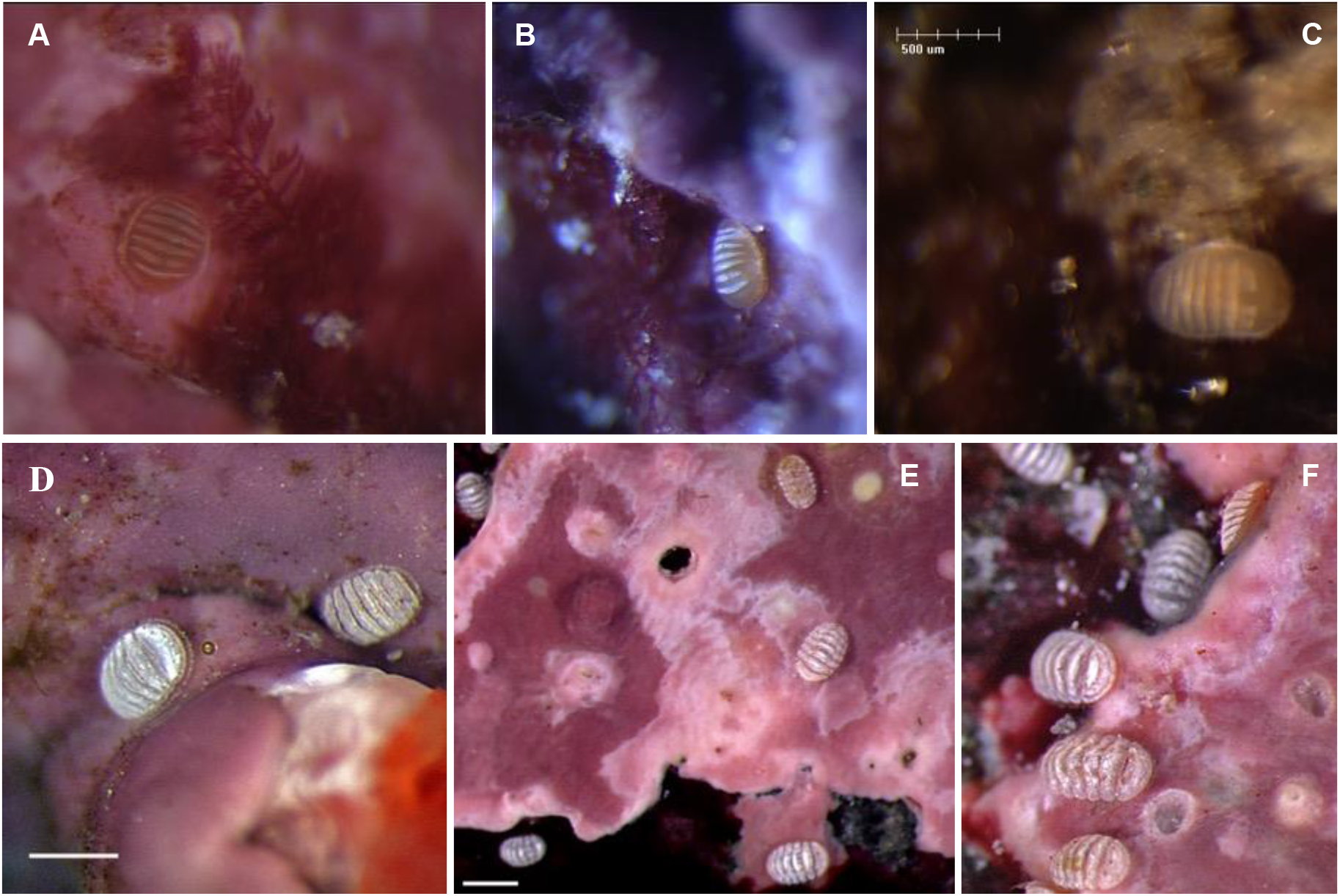
*Katharina tunicata* post-metamorphic juveniles on encrusting coralline red algae A, B) 1 dpm, C) 3 dpm, D) 14 dpm and E, F) 36 dpm.

**Figure 17.**
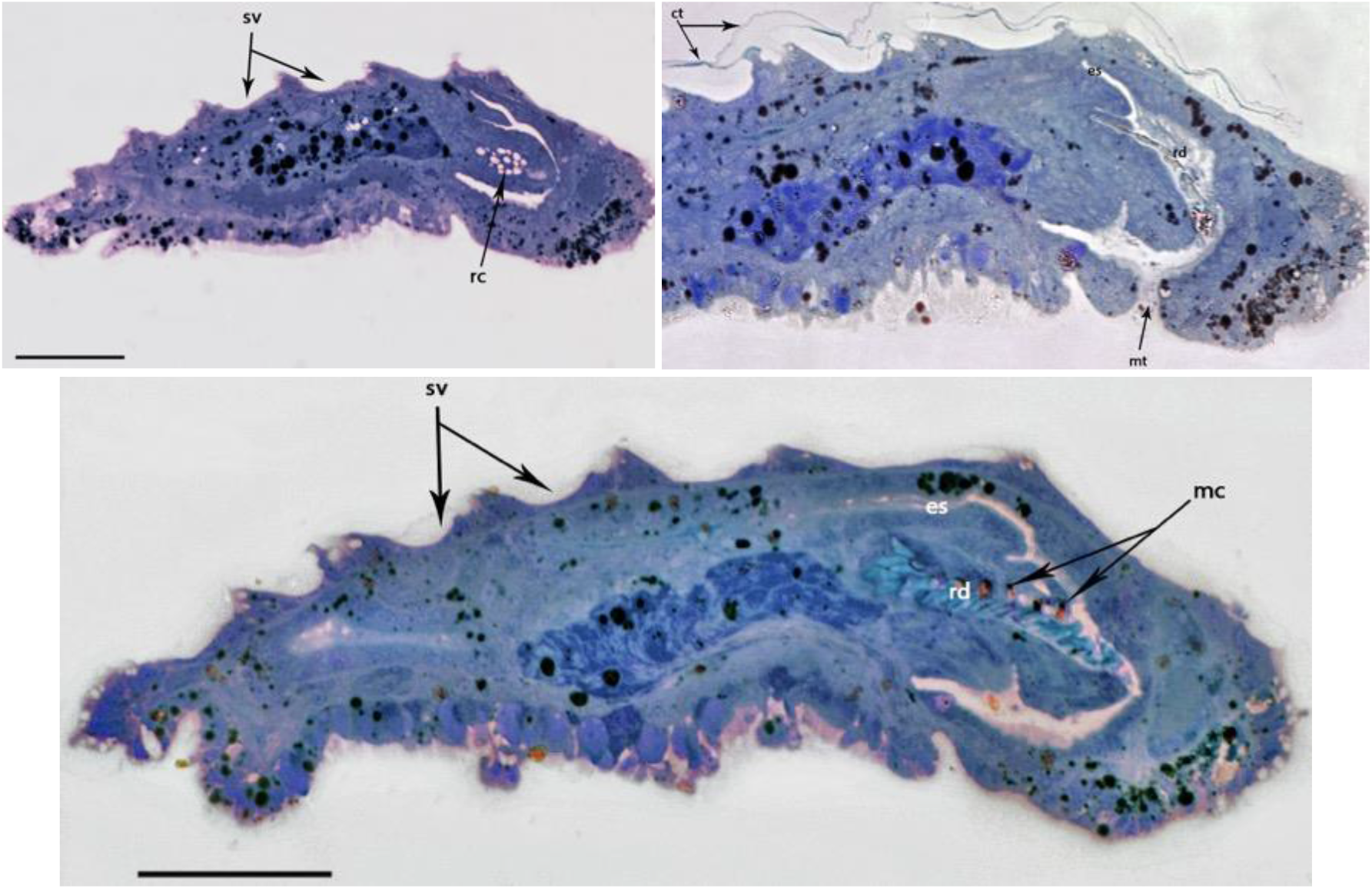
*K. tunicata* 14 dpm longitudinal histological sections 1 μm thick, tissue coloured with Richardson’s stain. Scales are representative of 50 μm; ct, cuticle; es, esophagus; int, intestine; m, mouth; mc, magnetite caps; pg, pedal glands; rc, radular cartilages; rd, radula; sv, shell valves (decalcified).

**Figure 18.**
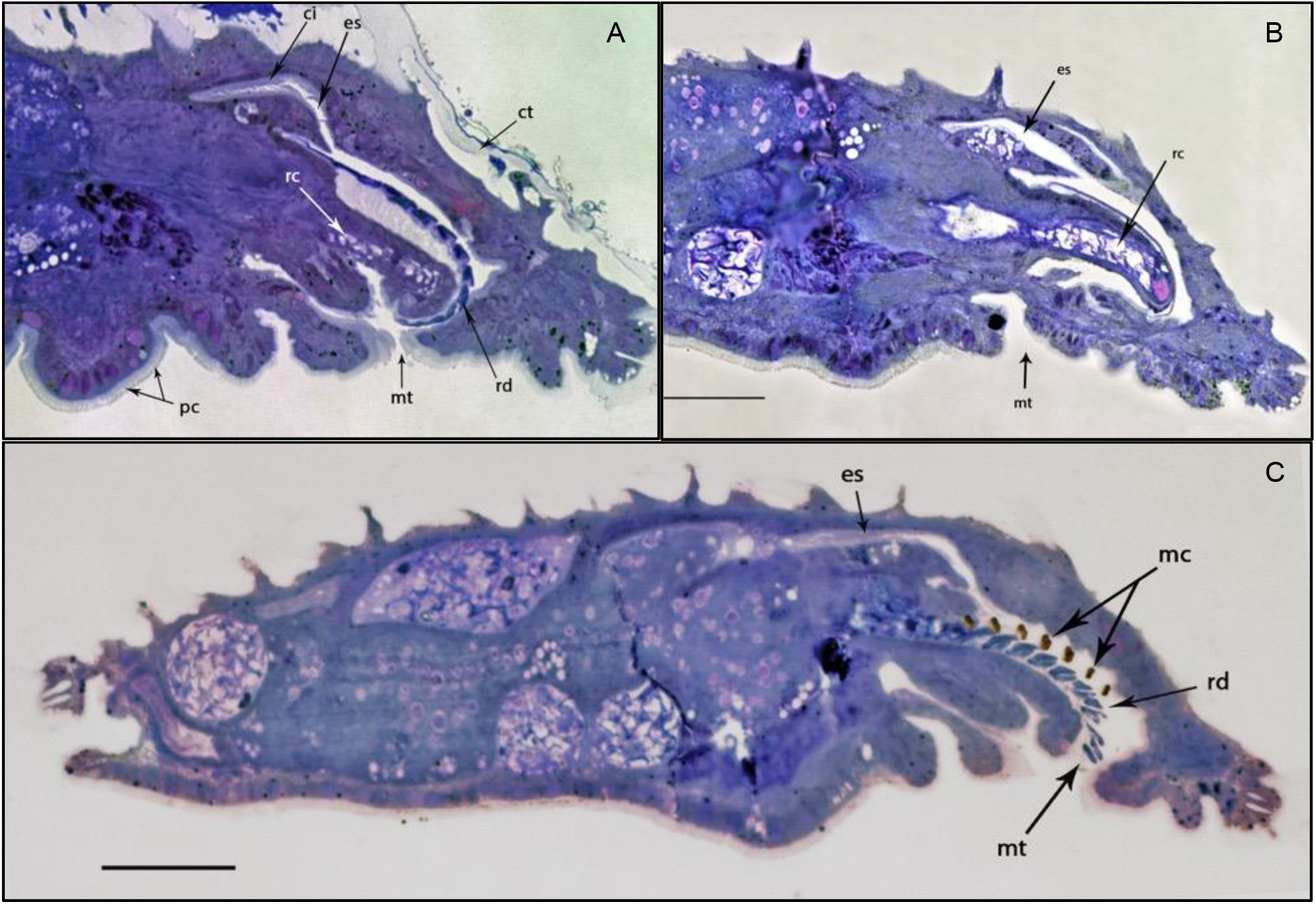
Longitudinal histological sections of 35-day post-metamorphic *Katharina tunicata* juveniles (1 μm), showing extensive foregut development. Scale bars represent 50 μm; ci, ciliated; ct, cuticle; es, esophagus; rc, radular cartilages; rd, radula; mc, magnetite caps; mt, mouth; pc, pedal cilia.

## Appendix II Tables

**Table 1.** Major events in the development of *K. tunicata* and whether they were observed from whole mount (wm) or thin section (ts) at time first observed relative to fertilization and hatching

| **Event** | **Specimen** | **Time**<br>(days past<br>fertilization) | **Time**<br>(days past hatching) |
| --- | --- | --- | --- |
| 8-cell stage | wm | 1 | – |
| Gastrulation | wm | 1.5 (12–36 h) | – |
| Hatching | wm | 2.5 (36–60 h) | – |
| Larval eyespots | wm | 6.5 | 4 |
| Spicules | wm | 10.5 | 8 |
| 6 Shell valves | wm | 10.5 | 8 |
| Buccal mass | ts | 12.5 | 10 |
| Esophagus | ts | 12.5 | 10 |
| Neuropil | ts | 12.5 | 10 |
| Radular rudiment | ts | 19.5 | 17 |
| Competent to metamorphose | wm / ts | 19.5 | 17 |
| Metamorphosis | – | 20.5 | 18 |
| 7 th shell valve | wm | 21.5 | 19 |
| Pallial groove | ts | 21.5 | 19 |
| Well-developed radula | ts | 34.5 | 32 |

**Table 2.** Comparison of developmental times reported in the literature for species of Polyplacophora. Times are all reported in days post fertilization.

| Citation | Hatch | Eyes | Spicules | Shell | Settle | Metamorphose |
| --- | --- | --- | --- | --- | --- | --- |
| Biggar, 2018, *K. tunicata* | 2.5 | 6.5 | 10.5 | 10.5 | 19.5 | 20.5 |
| Leise, 1984, *M. muscosa* | 1 | – | 5.5-6 | 10-13 | 9-10 | 12 |
| Lord, 2011, *C. stelleri* | 2 | 3.8 | – | – | 5 | – |
| Watanabe and cox, 1974,<br>*M. muscosa* and *M. lignosa* | 1 | 4 | 5 | 7-13 | 5.5-11.5 | 6.5-13.5 |

